# Alterations of the skeletal muscle nuclear proteome after acute exercise reveals a post-transcriptional influence

**DOI:** 10.1101/2024.08.08.607176

**Authors:** Ryan A. Martin, Mark R. Viggars, James A. Sanford, Zane W. Taylor, Joshua R. Hansen, Geremy C. Clair, Joshua N. Adkins, Collin M. Douglas, Karyn A. Esser

**Affiliations:** Department of Physiology and Aging, University of Florida, Gainesville, FL, USA; Myology Institute, University of Florida, Gainesville, FL, USA; Pacific Northwest National Laboratory, Richland, WA, USA; Department of Biomedical Engineering, Oregon Health and Science University, Portland, OR, USA

**Author notes:** Correspondence: Karyn A. Esser, Ph.D.

**Keywords:** Exercise, Muscle, Nuclear Proteome, Nuclei, Proteomics

## Abstract

Exercise is firmly established as a key contributor to overall well-being and is frequently employed as a therapeutic approach to mitigate various health conditions. One pivotal aspect of the impact of exercise lies in the systemic transcriptional response, which underpins its beneficial adaptations. While extensive research has been devoted to understanding the transcriptional response to exercise, our knowledge of the protein constituents of nuclear processes that accompany gene expression in skeletal muscle remains largely elusive. We hypothesize that alterations in the nuclear proteome following exercise hold vital clues for comprehending the transcriptional regulation and other related nuclear functions. We isolated skeletal muscle nuclei from C57BL/6 mice both sedentary control and one-hour post 30-minute treadmill running, to gain insights into the nuclear proteome after exercise. A substantial number of the 2,323 proteins identified, were related to nuclear functions. For instance, we found 59 proteins linked to nucleocytoplasmic transport were higher in sedentary mice compared to exercise, hinting at an exercise-induced modulation to nuclear trafficking. Furthermore, 135 proteins exhibited increased abundance after exercise (FDR < 0.1) while 89 proteins decreased, with the most prominent changes in proteins linked to mRNA processing and splicing. Super resolution microscopy further highlights potential localization change in mRNA processing proteins post-exercise, further suggesting changes in nuclear transport dynamics. Nonetheless, our data provide important considerations for the study of the nuclear proteome and supports a paradigm through which exercise downregulated mRNA processing and splicing, offering valuable insights into the broader landscape of the impact from acute exercise.

**New & Noteworthy:** Exercise plays a crucial role in promoting muscle health, but our understanding of nuclear proteins orchestrating exercise responses is limited. Isolation of skeletal muscle nuclei coupled with mass spectrometry enhanced the identification of nuclear proteins. This approach was used to investigate the effects of acute exercise, revealing changes in the muscle nuclear proteome 1-hour post-exercise, including proteins linked to post-transcriptional processing and splicing. Our findings offer insights into the exercise-induced changes within muscle nuclear proteins.

## Introduction

Exercise is widely acknowledged as a pivotal contributor to overall well-being and plays a crucial role in both preventing and treating various diseases. Its manifold benefits are extensively documented, as are the intricate signaling pathways that trigger exercise-induced changes in gene expression(1, 2). Although alterations in gene expression post-exercise are orchestrated within the nucleus, our comprehension of the responses at the level of nuclear proteins remains limited. Hence, the primary objective of this study is to offer a deeper understanding of the skeletal muscle nuclear proteome and its response to acute exercise.

Recent research has ventured into exploring nuclear processes that extend beyond transcription in skeletal muscle. This includes the influence of exercise on DNA methylation and chromatin modifications. Exercise often induces alterations in the methylation status of genomic regions, thereby affecting gene expression(3, 4). Concurrently, exercise-related signaling cascades, such as AMP-activated protein kinase (AMPK)(5, 6) and Ca2+/calmodulin-dependent kinase (CaMKII)(6, 7), can prompt chromatin remodeling through chromatin-targeting factors, thus modifying the epigenetic landscape. Despite these advancements, our knowledge of nuclear proteins orchestrating gene expression, inclusive of transcription, post-transcription, epigenetic, chromatin modeling, transcript transport, etc., in skeletal muscle, particularly following exercise, remains notably deficient. Therefore, a comprehensive exploration of nuclear proteins facilitating these modifications is warranted.

An inherent challenge in studying the nuclear proteome within large-scale proteomics lies in the limited coverage of small, less abundant proteins, including transcription factors and other proteins that reside in or attach to the nucleus. The intricacies of skeletal muscle add further complexities, as fundamental contractile proteins constitute a substantial proportion of the total protein mass(8), further exacerbating the dynamic range challenges for mass spectrometry-based proteomics. Consequently, strategies to address these challenges, such as subcellular fractionation, are frequently employed to investigate organelle-resident proteins(9, 10). Past research has demonstrated the ability of subcellular fractionation to isolate distinct organelle proteomes, including nuclei, mitochondria, Golgi apparatus, and the endoplasmic reticulum (ER)(11, 12). More recently, Cutler et al. applied a similar approach for the specific isolation of skeletal muscle nuclei in the context of muscle aging(13). However, such a strategy has not yet been employed to explore skeletal muscle nuclei in the context of exercise.

In this study, our aim is to define the skeletal muscle nuclear proteome’s response to acute exercise. Our approach initially involves the isolation of skeletal muscle nuclei to establish the nuclear proteome in the absence of exercise. Subsequently, we delve into the effects of acute exercise on the nuclear proteome, uncovering significant changes related to mRNA processing and splicing. In doing so, our data offers valuable insights into the nuclear proteome landscape of skeletal muscle following acute exercise.

## Materials and Methods

### Animals

All animal experimental procedures were approved and conducted under the guidelines of the University of Florida Institutional Animal Care and Use Committee (IACUC #201809136). The use of animals for exercise protocols was in accordance with guidelines established by the US Public Health Service Policy on Humane Care and Use of Laboratory Animals. Male C57BL/6 mice (Charles River) were single housed with food and water provided *ad libitum* with experimental intervention and tissue collected at 5 months of age (n = 20).

### Exercise Familiarization and Testing

Prior to maximal treadmill testing, animals were exposed to 3 familiarization bouts with the treadmill. For maximal treadmill testing, following a 5-minute warm-up at 10 cm/s at a 10° incline, treadmill speed increased by 3 cm/s at 15° incline every 2 minutes until visible exhaustion. Work done was calculated using the following formula: Work Done = Mass * Distance * sinθ. Mice then performed an acute bout of treadmill running (n = 10) at 70% work done for 30 minutes at 15° incline while unexercised controls remained sedentary (n = 10), with tissues collected in both groups 1-hour post-exercise (adapted from (14)). Maximal exercise testing results for sedentary and exercise mice are provided in supplementary figure 1 (**Supplemental Figure S1**).

### Body Composition

Body composition was determined prior to the experimental procedures and exercise on all animals by magnetic resonance imaging use an EchoMRI. The Imaging system was calibrated with standard operating procedures prior to testing.

### Nuclei Isolation

Frozen whole gastrocnemius muscles (4 pooled gastrocnemius muscles) were placed in Homogenization Buffer (10 mM HEPES, pH 7.5, 10 mM MgCl_2_, 60 mM KCl, 300 mM Sucrose, 0.1 mM EDTA, and complete mini protease inhibitor (Millipore; Cat. #11836170001) and homogenized with a Polytron homogenizer. Homogenates were centrifuged for 3 minutes and 17 x g at 4° C and the supernatant were collected. An additional 2 mL of homogenization buffer was added to the pellet followed by centrifugation at 17 x g for 3 minutes at 4° C. This process was repeated 4 times. Pooled supernatants were filtered by 40 µm filters by a simple pour and repeated for a total of 3 filtrations. The filtrate was centrifuged at 17 x g for 10 minutes at 4° C followed by collecting the supernatant and centrifuged at 1000 x g for 10 minutes at 4° C. After discarding the supernatant, pellets were resuspended in homogenization buffer and rotated at 4 °C for 10 minutes. ∼200 µL of sample was then placed on top of 2.15 M Sucrose cushion and centrifuged at 13,500 x g at 4° C for 90 minutes. Following the centrifuging of samples, a small volume of the top layer of sucrose cushions were removed by vacuum suction followed by the addition of ∼200 µL of homogenization buffer. This small volume of the topmost layer of sucrose cushions and homogenization buffer wash was repeated three times to remove potential debris remaining on top of the sucrose cushion. Entire sucrose cushions were then removed and pelleted (nuclei) fractions were resuspended in homogenization buffer and combined for each sample and centrifuged at 1,000 x g for 5 minutes at 4° C. The supernatants were removed, and the nuclei pellets were stored at −80° C until analysis. In total, 4 muscles were pooled (i.e., 2 mice) per 1 sample for proteomic analysis, resulting in 4 samples for sedentary controls (n = 4) and 4 samples for exercise (n = 4), identified as ‘post-sucrose’. Additional samples were included for comparison prior to nuclei isolation (n = 1) for proteomic analysis identified as ‘pre-sucrose’.

### Immunohistochemistry

Isolated nuclear fraction prior to and post-sucrose cushions were used for immunohistochemical detection of nuclei and cytoplasmic debris. Nuclear fractions were mixed with Phalloidin conjugated AlexaFluor™ 488 (Invitrogen Cat. #A12379; 1:1,000) to identify cytoplasmic F-actin fragments and 4’,6-diamidino-2-phenylidole (DAPI) (Invitrogen Cat. #D1306; 1:1,000) to label nuclei. Labeled fractions were visualized with a NIKON Ti2 inverted epifluorescence microscope.

### Super-resolution Microscopy

As previously described (15), 10μm thick longitudinal cryosections of tibialis anterior muscle were allowed to equilibrate from −80°C to room temperature for 15 minutes. Individual cryosections were rehydrated using Tris-Buffered Saline (TBS) and permeabilized with 0.5% Triton X-100 in TBS. Sections were washed TBS before a 30-minute incubation with Image-iT FX Signal Enhancer (Invitrogen). Following this incubation, sections were blocked for 1-hour using blocking buffer (5% Normal Goat Serum, 5% Bovine Serum Albumin, 5% Normal Alpaca Serum, 0.1% Triton X-100 in TBS). Sections were then incubated overnight at 4°C with primary antibodies (1:500) in antibody dilution buffer ((5% Bovine Serum Albumin in TBS-Tween (0.1%)). The following day, sections were washed with TBS-T solution, and once with TBS for five minutes. Sections were incubated with secondary antibodies (1:1000) in antibody dilution buffer for 1-hour at room temperature. Sections were then washed with TBS-T, followed by TBS prior to a 10-minute incubation of DAPI (1:10,000) in TBS. Sections were washed twice for five minutes with TBS, before treatment with TrueBlack for 30 seconds. Sections were washed twice with TBS, followed by a single 5-minute wash using ultrapure water. Slides were then mounted and cover-slipped using Prolong Glass antifade mounting media and allowed to cure overnight in the dark at room temperature. Sections were imaged on a Nikon Ti2-E inverted microscope equipped with a 100x oil-immersion objective (CFI Apochromat TIRF, NA 1.49) and a confocal spinning disk (Yokogawa, CSU-X1) with super-resolution imaging module (Gataca LiveSR Systems) and sCMOS camera (Teledyne Photometrics, Prime95B). Z-stacks were obtained in 0.1 µm increments set tp 500 ms exposure per individual channel and reconstructed using maximum intensity projections. Image analysis and processing were all performed using ImageJ software.

### Proteomics sample preparation

Isolated nuclei were prepped for LC-MS/MS based on a protocol described previously (16, 17). Briefly, nuclei pellets were lysed by adding 10 uL of 0.05% DDM in 50 mM Tris, pH 8.0 and incubating on a thermomixer set to 300 rpm and 37°C for 30 minutes, and protein concentrations were determined via BCA assay. Proteins were reduced with dithiothreitol (DTT, final concentration 5 mM) for 30 minutes at 37°C and alkylated with iodoacetamide (IAA, final concentration 10 mM) for 30 minutes at 25°C. Trypsin was added at a 1:50 enzyme:substrate ratio (100 ng trypsin added for samples below the BCA assay detection limit) and samples were digested for 2 hours on a thermomixer set to 300 rpm and 25°C. After 2 hours, additional trypsin was added (again at a 1:50 ratio, or 100 ng total for samples below BCA assay detection limit) and samples were digested for 14 hours on a thermomixer set to 300 rpm and 25°C. Digestions were quenched by adding formic acid to a final concentration of 1%. Samples were diluted to 30 uL with water, peptide concentration was determined via BCA assay, and peptides were vialed at 0.1 ug/uL in plastic vials coated with 0.01% DDM for LC-MS/MS analysis.

### LC-MS/MS Analysis

5 uL of the 0.1 ug/uL digested peptides were analyzed by reverse-phase LC-MS/MS using a Dionex Ultimate 3000 RSLC nanopump (ThermoFisher Scientific) coupled with a QExactive HF mass spectrometer (ThermoFisher Scientific). The LC was configured to first load the sample on a solid-phase extraction (SPE) column (150 um inner diameter, packed with 5 um Jupiter C18 material (Phenomenex) followed by separation over a 120-minute gradient on an analytical column (25 cm, 75 um inner diameter, packed with 1.7 um BEH C18 material (Waters). Effluents were analyzed with the Orbitrap mass spectrometer operated in the data-dependent acquisition mode, with the top 12 ions from survey scans selected for high-energy dissociation. An isolation window of 0.7 Da was used for the isolation of ions, and a collision energy of 30% was used for high-energy collisional dissociation with an automatic gain control setting of 3×10^6^ ions. MS/MS scans were acquired at a resolution of 30,000 with an AGC setting of 1×10^5^ ions and a maximum injection time of 100 ms. Mass spectra were recorded for 120 min with a dynamic exclusion window of 45 seconds.

### LC-MS/MS Data Processing

Raw LC-MS/MS datasets were searched using MaxQuant software (v1.6.12.0) against a mouse UniProt database downloaded in January 2021, with match-between-run (MBR) and iBAQ calculations enabled. The following MaxQuant settings were used: tryptic digest with a maximum of 2 missed cleavages; fixed modification of cysteine carbamidomethylation; variable modifications of methionine oxidation and n-terminal acetylation; 0.7-minute matching window frame. Protein abundances detected by LC-MS were used for fold change calculations in appropriate figures. Prior to statistical analysis, protein intensities were log2-transformed, median-centered within samples, and data was filtered to unique proteins identified in at least 50% of samples. Statistical analyses were performed in RStudio software using moderated t-tests from the *limma* package.

### Statistical Analysis

Data sets were analyzed for normality and equal variance prior to statistical inference testing. T-tests were used between sedentary and exercise mice body composition and exercise performance using Prism Graphpad 9.1.2 for Windows (San Diego, California, USA). Prior to statistical inference testing, *a priori* P < 0.05 was used to determine statistical significance. T-test data sets are reported as Mean ± SEM. Statistical significance within proteomic datasets was determined using a false discovery rate (FDR < 0.1). Functional cluster analysis was performed using the Database for Annotation, Visualization and Integrated Discovery (DAVID) (18, 19) in addition to Gene Set Enrichment Analysis (GSEA) (20, 21) for significantly enriched (FDR < 0.05) Biological Processes, Cellular Compartments, Reactome, and KEGG pathways on the Mus Musculus genome background.

## Results

### Experimental Approach and Animal Performance

The general research design overview is provided in Figure 1A. We obtained body composition measurements on all mice in this study prior to the maximal exercise test and found that all mice were similar in body mass (**Figure 1B**) (27.3 ± 0.77 g Sed vs 27.31 ± 0.43 g Ex; P > 0.05) as well as body composition, including lean mass (22.1 ± 0.44 g Sed vs. 22.4 ± 0.27 g Ex) (**Figure 1C**) and fat mass (3.69 ± 0.47 g Sed vs. 3.33 ± 0.30 g Ex; P > 0.05) (**Figure 1D**). To appropriately define the relative exercise bout for the study, all mice performed a progressive maximal treadmill test demonstrating similar exercise capacities as presented in supplemental figure 1 (**Supplement Figure S1**). Seven days following the maximal treadmill test, mice were divided into either sedentary (n = 10) or exercise (n = 10) groups with exercise mice performing an acute bout of exercise consisting of 70% of work done. The acute exercise bout was performed at an average speed of 26.53 ± 0.51 cm/s (**Figure 1E**) with total distance of 478 ± 2 m (**Figure 1F**) and a total of 3384.9 ± 6.35 A.U. of work done (**Figure 1G**) by the mice of the exercise group.

**Figure 1.**
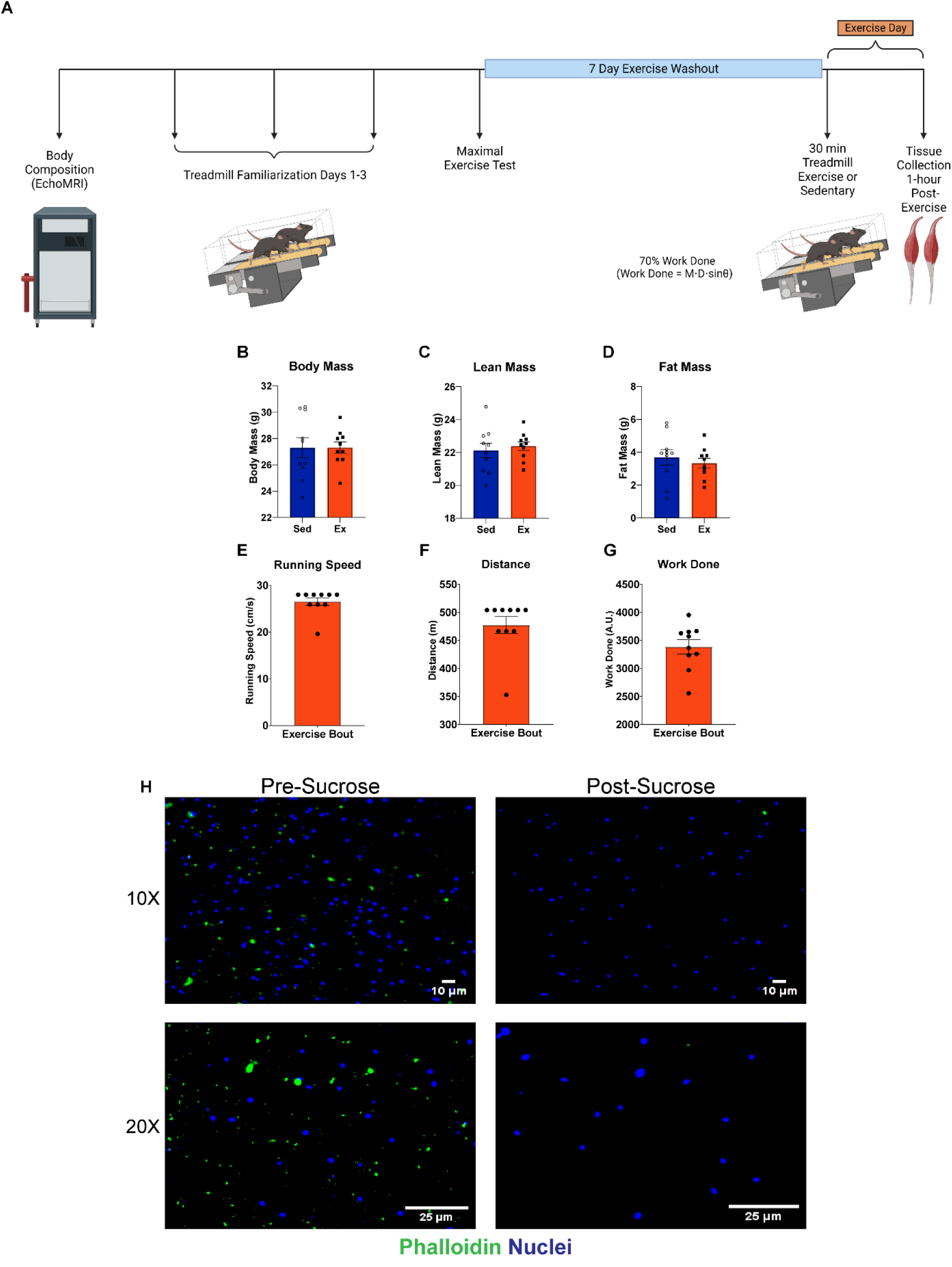
Experimental Approach and Phenotypic and Performance of Sedentary and Exercise Mice. **A)** Experimental timeline including Body Composition measurements, treadmill familiarization and the maximal exercise test. Following a 7-day exercise washout period, mice performed an acute bout (30 minutes) of treadmill running at 70% work done (Work done = mass * distance * sinθ) and muscles collected 1-hour post-exercise. **B-D)** Body composition measurements for body mass (B), absolute (C) lean and fat (D) mass indicating similar phenotypes between sedentary and exercise mice. Independent t-tests between Sed and Ex groups; all measures were not significant (NS; P > 0.05). **E-G)** Similar Performance measures of the acute exercise bout for running speed (E), distance (F), and work done (G). **H)** Immunohistochemistry images of isolated skeletal muscle nuclei (DAPI; blue) and cytoplasmic debris (Phalloidin; green) prior to (*left panels*) and after (*right panels*) the sucrose cushion. Top and bottom panels are taken with 10X (Scale bar = 10 µm) and 20X (Scale bar = 25 µm) objectives, respectively. Panel A was created using BioRender.com.

In order to investigate the nuclear proteome, we isolated skeletal muscle nuclei using a modified isolation protocol previously described(22, 23). Following homogenization of gastrocnemius muscle tissue, lysates were passed through a 40 µm filter to remove large aggregates of debris and subjected to high-speed centrifugation on a sucrose cushion for the isolation of nuclei. As a qualitative measure of nuclear enrichment, homogenates were immunolabelled with Phalloidin (*green*) to identify any cellular debris (e.g., actin cytoskeleton) and DAPI (*blue*) for nuclei before and after the sucrose cushion (**Figure 1H**). While we observed a large population of nuclei after bulk homogenization, there remained a substantial abundance of phalloidin^+^ debris in homogenates prior to the sucrose cushion (**Figure 1H, *left panels***). However, limited phalloidin^+^ debris was then found in homogenates after the sucrose cushion (**Figure 1H, *right panels***), indicating the reduction of cellular debris and cytoplasmic material in the remaining population of nuclei. Importantly, since whole muscle was used for the isolation of nuclei, we cannot rule out the incorporation of nuclei from cell types other than skeletal muscle (e.g., endothelial, immune, nerve). Research has estimated upwards of ∼70-90% of nuclei in whole muscle homogenates are skeletal muscle nuclei residing within the myofiber(13, 24, 25). Nonetheless, here we show the successful isolation of skeletal muscle nuclei between phenotypically similar animals to determine changes within the nuclear proteome after acute exercise.

### The Skeletal Muscle Nuclear Proteome

Our initial objective was to characterize the nuclear proteome of skeletal muscle using a fractionation technique involving a sucrose cushion which yielded a purer fraction of nuclei (**Figure 1**). To assess the level of protein enrichment we performed proteomics on the filtered fraction before sucrose (referred to as “pre-sucrose”; n = 1), as well as isolated nuclei after fractionation (termed “post-sucrose”; n = 4) from the gastrocnemius muscles of sedentary mice. To extend our analysis in this section we also compared our results to those published for whole mouse muscle(26).

We first compared proteins between whole muscle, “pre-sucrose”, and “post-sucrose” datasets to identify commonly found proteins regardless of experimental approach. In this comparison, 620 proteins were shared among all three datasets (**Figure 2A**). Functional enrichment analysis (FDR < 0.05) unveiled a significant number of proteins related to mRNA processing (n=79) and splicing (n=71), and ribosome (n=50) (**Figure 2B and 2C**). As expected, some proteins more common to the cytoplasm including electron transport (n=31), glycolysis (n=9), and cardiac muscle contraction (n=23) were detected, underscoring the combination of their relative abundance and potential carry over into the post-sucrose fraction.

**Figure 2.**
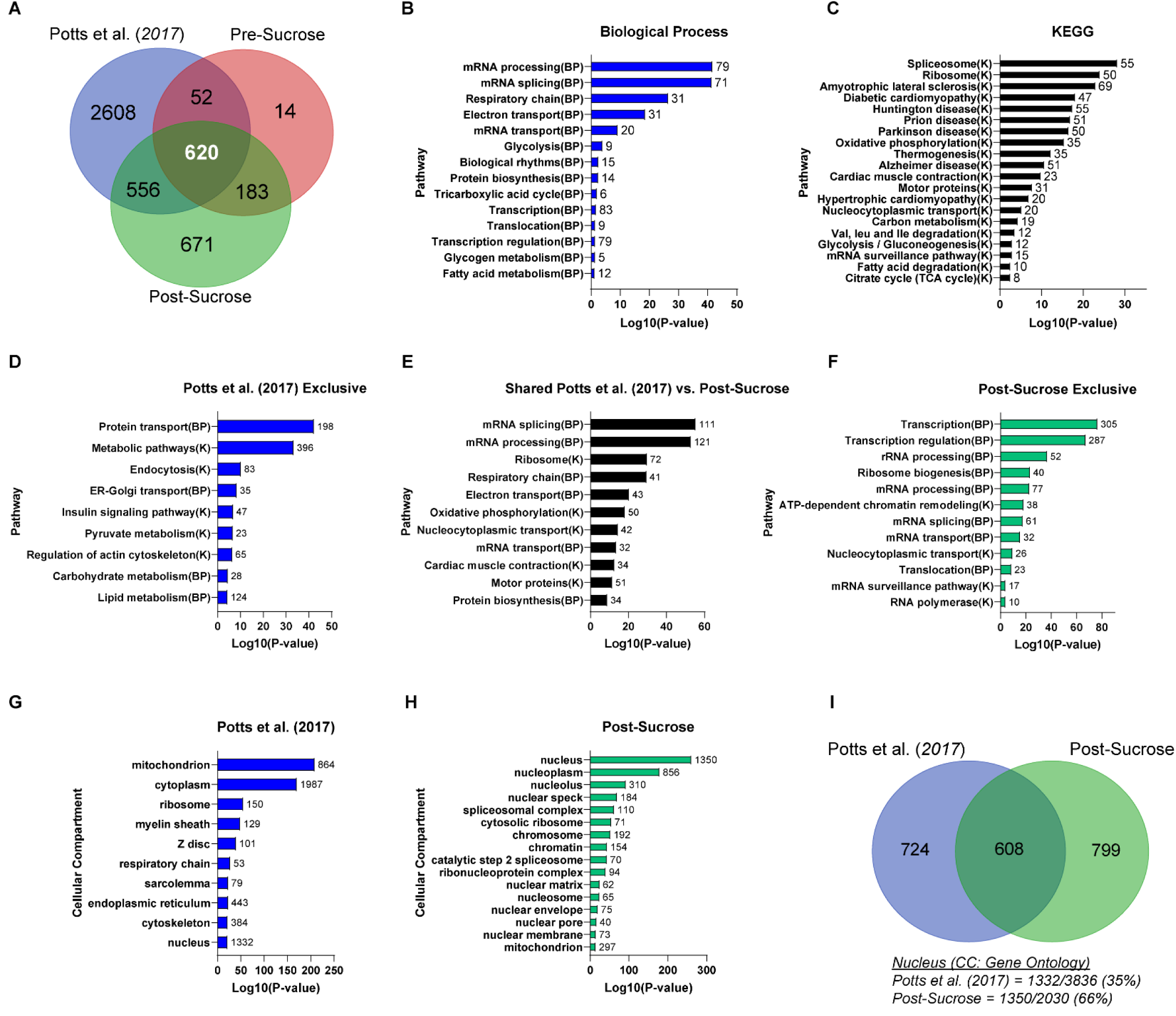
Skeletal Muscle Nuclei Proteomics. **A)** Venn diagram of proteins found in whole muscle homogenate (*blue*) (Potts et al. 2017), pre-sucrose (*red*), and post-sucrose (*green*). **B-C)** Pathway enrichment for Gene Ontology biological processes (B; *blue bars*) and KEGG (C; *black bars*) of the 620 common proteins between all comparisons in A. **D-F)** Pathway enrichment analysis using biological processes and KEGG of proteins unique to whole muscle (D), shared between whole muscle and “post-sucrose” nuclei (E), and proteins found exclusively in the “post-sucrose” nuclei (F). Pathways are also noted as biological process (BP) or KEGG (K) with the number of proteins in each pathway beside the respective bar. **G-H)** Gene Ontology analysis for cellular compartment of proteins identified in whole muscle (G) and “post-sucrose nuclei (H), with the number of proteins enriched in each compartment beside each bar. **I)** Venn diagram of proteins enriched in “Nucleus” compartment from whole muscle (*blue*) and “post-sucrose” nuclei (*green*) in G and H. Relative percentages of nuclear compartment enriching proteins in whole muscle (35%) and “post-sucrose” nuclei (66%) are displayed below venn diagram.

From our proteomics analysis, we detected 869 proteins in the pre-sucrose fractions and 2,030 proteins in the post-sucrose fraction. Of the 869 pre-sucrose proteins, 803 (92%) were shared with the post-sucrose fraction. The post-sucrose fraction yielded 2,030 proteins, of which 1,227 were not identified in the pre-sucrose fraction. These findings highlight the extended protein identification provided by the inclusion of the sucrose cushion step for the subsequent enrichment of the nuclear associated proteome. Since we found fewer proteins uniquely identified and quantitated in the “pre-sucrose” fraction, we focused our analysis between “post-sucrose” nuclei and whole muscle.

We next compared the results from the “post-sucrose” fraction to the whole muscle proteome to provide insight into the ability to identify more nuclei associated proteins. For this comparison there were 3,836 proteins in whole muscle homogenate compared to the 2,030 proteins in the post-sucrose fraction. Importantly, we found 1,176 shared proteins between these two datasets, with 2,660 proteins unique to whole muscle and 854 proteins unique to the “post-sucrose” nuclei. The number of unique post-sucrose proteins suggests that we were successful enriching for nuclei associated proteins that likely remained undetected when performing proteomics from whole muscle lysates. Functional enrichment of the whole muscle unique 2,660 proteins revealed an abundance of proteins related to protein transport (n=198), metabolic pathways (n=396), signaling pathways (n=47), and the actin cytoskeleton (n=65) (**Figure 2D**). The 1,176 shared proteins between whole muscle and “post-sucrose” nuclei enriched for some nuclear processes like mRNA splicing (n=111) and mRNA processing (n=121) as well as inclusion of ribosome (n=72), electron transport (n=43), and cardiac muscle contraction (n=34) (**Figure 2E**). For the 854 proteins exclusively identified in our “post-sucrose” nuclei, we see the emergence of unique nuclear centric processes including transcription (n=305), followed by rRNA processing (n=52), chromatin remodeling (n=38), and mRNA transport (n=32) (**Figure 2F**). These results confirm that the sucrose fractionation successfully isolates nuclei from our skeletal muscle samples, allowing for deeper coverage of the nuclear proteome.

We next used the Gene Ontology: Cellular Compartment database for analysis of the whole muscle and our “post-sucrose” nuclei. Some of the most significantly enriched (FDR < 0.05) cellular compartments in whole muscle were mitochondrion, cytoplasm, ribosome, and Z disc (related to contractile proteins) (**Figure 2G**). However, 1,332 proteins that are annotated to the nucleus were identified in the whole muscle fraction. Comparatively, the post-sucrose unique proteins largely enriched for sub-nuclear compartments, nucleolus, splicesomal complex, nuclear envelope and chromatin (FDR < 0.05) (**Figure 2H**). Comparing the number of proteins enriching for the nucleus cellular compartment (1,332 whole muscle vs. 1,350 “post-sucrose”), we found 608 proteins are shared with 799 proteins exclusively identified in our “post-sucrose” nuclei (**Figure 2I**). Importantly, out of the total number of proteins identified in either proteome, 35% of proteins within whole muscle were identified as part of the nuclear compartment, compared to the 66% of proteins in our “post-sucrose” nuclei.

Collectively, these results demonstrate the ability to significantly improve mass spectrometry-based identification of nuclear proteins in skeletal muscle tissue through the utilization of a fractionation approach compared to conventional whole muscle proteomics. The heightened resolution of nuclear resident proteins in skeletal muscle provides opportunities for more nuanced investigations into alterations within the nuclear proteome in response to diverse stimuli.

### Identified Transcription Factors in Isolated Muscle Nuclei

One of the inherent challenges in large-scale proteomics in skeletal muscle is the study of transcription factors. Based on our enrichment for nuclear proteins, we extended our exploration of proteins that might be involved in nuclear-specific functions, such as transcription, by utilizing the Gene Ontology: Molecular Function database (FDR < 0.05; **Figure 3A**). Using proteins enriched in the nuclear compartment of whole muscle and “post-sucrose” nuclei (**Figure 2I**), we highlight the distribution of proteins with relevant nuclear functions, such as DNA binding, RNA binding, chromatin binding, and transcription. Out of the total number of proteins identified in the respective proteomes, proteins related to DNA Binding (4.8%), RNA Binding (6.9%), Chromatin Binding (2.5%), and mRNA Binding (2.7%) from Potts et al.(26) covered <7% of the total number of proteins identified (n = 3,836). In contrast, “post-sucrose” nuclei contained a substantially higher proportion of proteins linked to DNA Binding (18.8%), RNA Binding (15.7%), Chromatin Binding (8.5%), and mRNA Binding (5.3%). Furthermore, the number of proteins in DNA Binding (n = 266), chromatin binding (n = 110), and transcription-related pathways like helicase activity (n = 66), coactivator activity (n = 46), and corepressor activity (n = 34) were exclusively found in the “post-sucrose” nuclei (**Figure 3B**), extending the number of nuclear function proteins compared to whole muscle.

**Figure 3.**
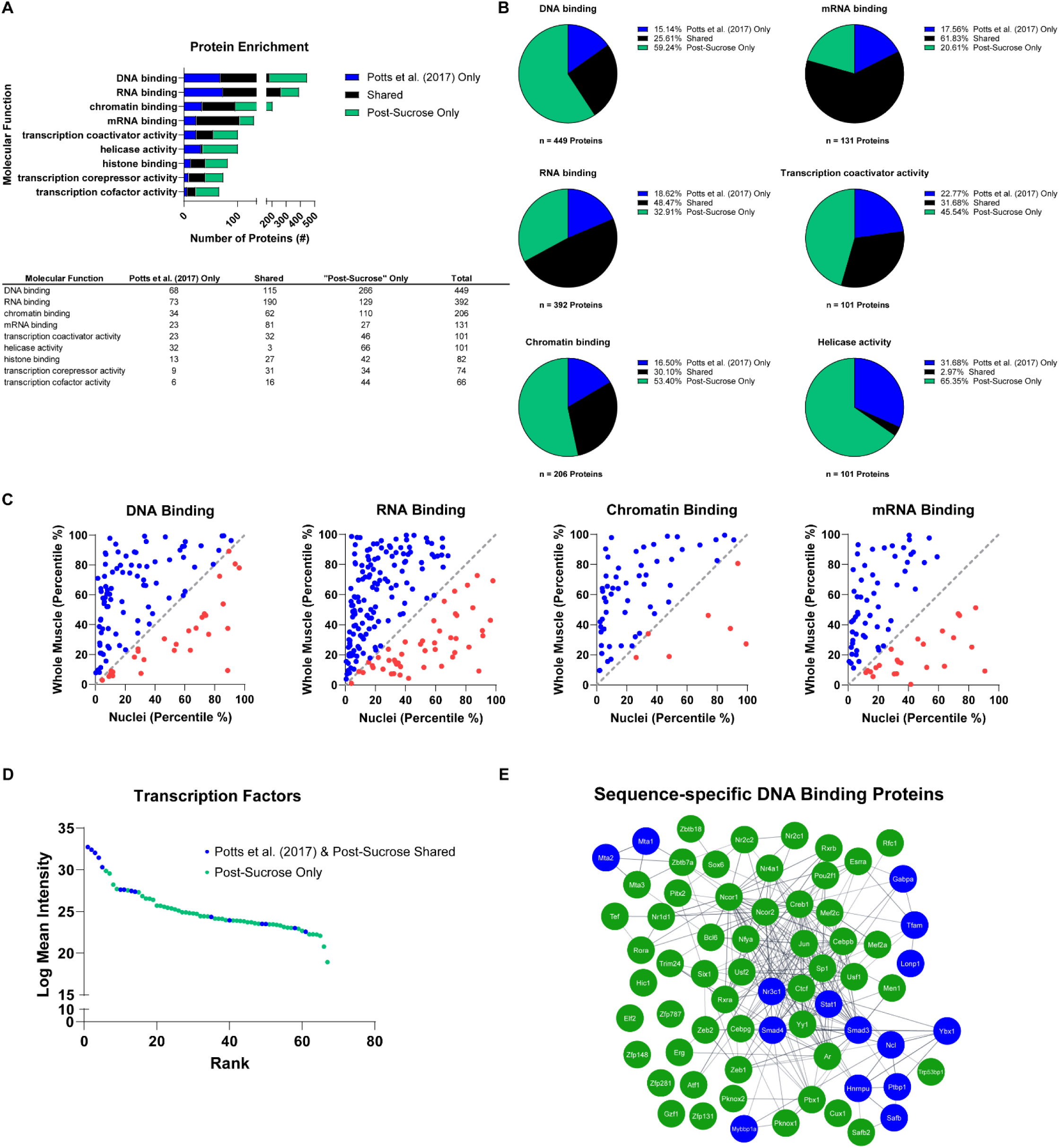
Transcription Factors in Skeletal Muscle Nuclear Proteome. **A)** Gene Ontology analysis for molecular function (FDR < 0.05) on nuclear compartment enriched proteins (from Figure 2I) with the total number of proteins in each pathway. Respective number of proteins within each pathway are shown as those exclusively identified in whole muscle (*blue*), shared between whole muscle and “post-sucrose” nuclei (*black*), and exclusively identified in “post-sucrose” nuclei (*green*). Table includes number of proteins identified enriched in molecular function pathways. **B)** Pie graphs for the top 6 molecular functions from Gene Ontology Analysis in Figure 3A and percent distribution of proteins in whole muscle (*blue*), shared between whole muscle and “post-sucrose” nuclei (*black*), and exclusively identified in “post-sucrose” nuclei (*green*). Total number of proteins enriched in each pathway is noted below each pie graph. **C)** Correlational plots of shared proteins only within DNA binding, RNA Binding, Chromatin Binding, and mRNA Binding molecular function pathways in 3A. Correlational plots are shown for each molecular function with the inclusion of a line of identity (*gray*). Proteins were ranked based on their log mean intensity and normalized to the total number of proteins in their respective proteomes represented in percentiles (%) with those ranked higher in nuclei (*blue*) and those ranked higher in whole muscle (*red*). **D)** Ranked order of identified sequence-specific DNA Binding proteins (i.e., Transcription Factors) based on their log mean intensity. Graph includes factors shared between whole muscle and “post-sucrose” nuclei (*blue*) and factors exclusively identified in “post-sucrose” nuclei (*green*). **E)** Protein network of sequence-specific DNA binding proteins in D, with factors shared between whole muscle and “post-sucrose” nuclei (*blue*) and factors exclusively identified in “post-sucrose” nuclei (*green*).

Next, to investigate the relative enrichment of the nuclear proteins identified in both the whole muscle vs. “post-sucrose” nuclear fraction we performed a rank ordered analysis for DNA Binding, RNA Binding, Chromatin Binding, and mRNA Binding. For this we ranked shared proteins from respective proteomes based on their quantitative relative abundance derived from label-free proteomics. The positional ranking of proteins was then normalized to the total number of proteins in their respective proteomes (3,836 whole muscle vs. 2,030 “post-sucrose” nuclei) and presented in percentiles. Therefore, the same identified protein in the top 10% of “post-sucrose” nuclei and 90% of whole muscle demonstrates an enhanced ability to identify the protein in isolated nuclei. Most of the shared proteins between whole muscle and “post-sucrose” nuclei were at a lower percentile (*blue*) compared to the same protein in whole muscle; whereas fewer proteins remained at a higher percentile in whole muscle compared to “post-sucrose” nuclei (*red*) (**Figure 3C**). These findings demonstrate that proteins were more prominently identified in “post-sucrose” nuclei in comparison to the whole muscle proteome.

To delve deeper into proteins related to DNA binding, particularly transcription factors, we identified a total of 67 proteins enriched under the Gene Ontology term sequence-specific DNA Binding within our “post-sucrose” nuclei (**Figure 3C and 3D**). Notably, 15 transcription factor proteins were shared between the whole muscle dataset and “post-sucrose,” and an additional 52 proteins were exclusively found in the “post-sucrose” nuclei. Importantly, many of the factors shared between the whole muscle dataset and “post-sucrose” were among the highly abundant factors (**Figure 3D**). Those transcription factors found exclusively in the “post-sucrose” nuclei included many well known in the muscle field such as AR, SOX6, and CREB1(**Figure 3E**). These findings demonstrate increased capacity for detection of transcription factors in muscle and establish the platform to use this approach to begin to study change in response to muscle use.

### The Nuclear Proteome in response to an acute bout of exercise

In order to investigate how an acute exercise session affects the nuclear proteome, we conducted an analysis of nuclear proteomes from sedentary (n = 4) and exercised (1-hour post-exercise; n = 4) mice. Of the 2,323 total proteins identified in any of the nuclear isolations, we observed that 367 proteins were exclusively detected in the sedentary group, while 284 proteins were only detected in the exercised mice (**Figure 4A**).

**Figure 4.**
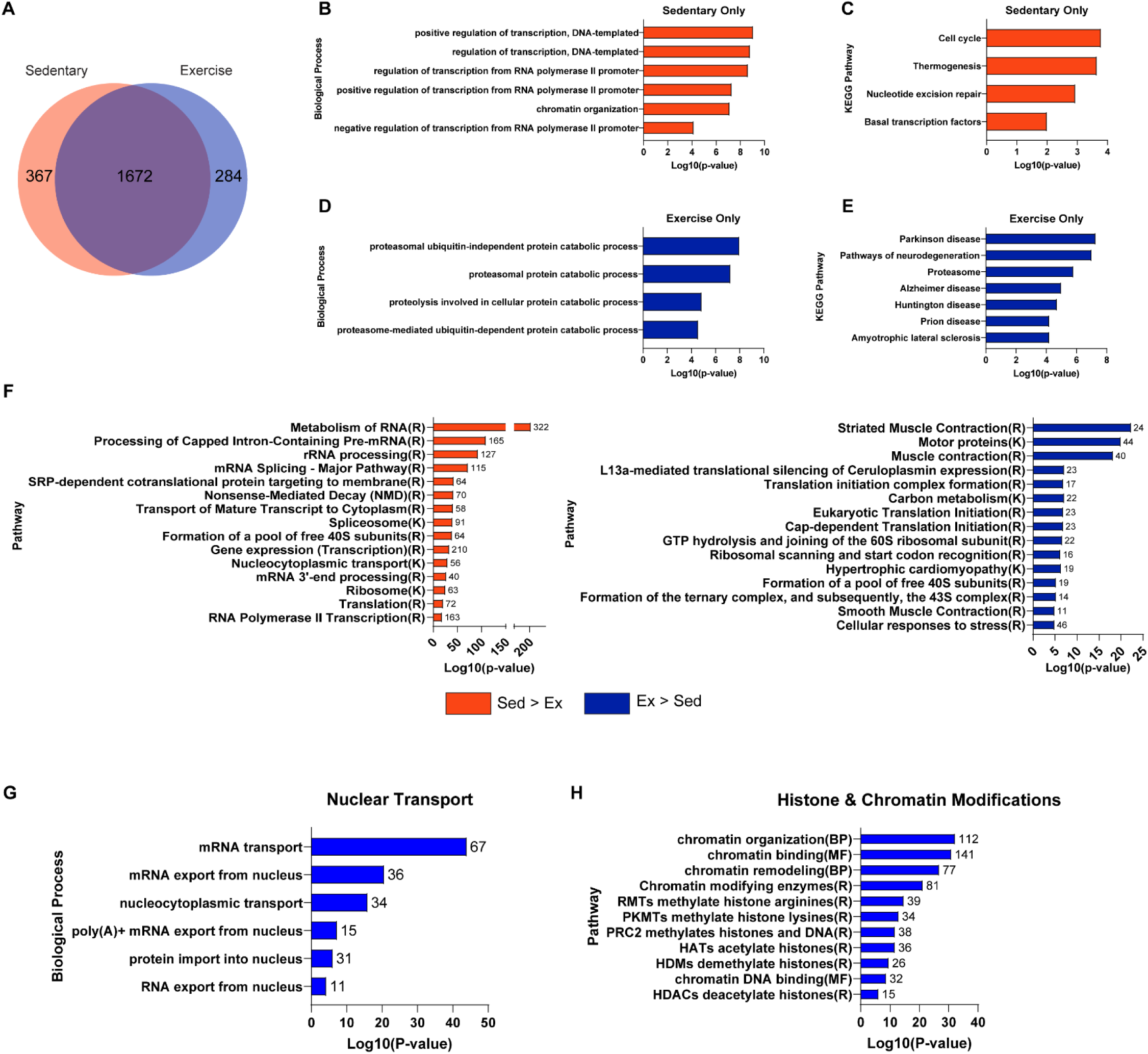
Comparison of sedentary and exercise mice nuclear proteomes. **A)** Venn diagram of proteins in isolated skeletal muscle nuclei from sedentary (*orange*) and exercise (*blue*) mice. **B-C)** Biological processes (B) and KEGG pathways (C) of proteins unique in sedentary muscle nuclei (367; *orange*). **D-E)** Biological processes (D) and KEGG pathways (E) of proteins unique in exercise muscle nuclei (284, *blue*). **F)** Top KEGG and Reactome pathways (FDR < 0.05) of shared proteins (1,672) with a FC > 1 higher in sedentary (*orange bars; 1,130*) or FC > 1 higher in exercise (*blue bars; 542*) mice. **G and H)** Pathway enrichments for proteins shared between sedentary and exercise nuclei related to the Biological Process ‘Nuclear Transport’ (G) and pathways related to histone and chromatin modifications (H). Numbers associated with bars reflect the number of proteins within a given pathway identified in our dataset. K = KEGG; R = Reactome; MF = Molecular Function.

Functional pathway analysis of the 367 proteins exclusive to sedentary mice revealed enriched biological processes related to transcription regulation, chromatin organization coupled with cell cycle regulation, thermogenesis, nucleotide excision repair, and basal transcription factors KEGG pathways (FDR < 0.05; **Figure 4B and 4C**). In contrast, the 284 proteins exclusively found in the skeletal muscle nuclei of exercised mice enriched for pathways in catabolic processes, proteasome-dependent mechanisms, and proteasomal subunit components (FDR < 0.05; **Figure 4D**). KEGG pathways enriched for the Proteasome pathway and several neurodegenerative diseases, including Parkinson’s, Huntington’s, and Alzheimer’s disease, which have previously been linked to proteasomal-related mechanisms(27–30) (**Figure 4E**). These findings suggest that, while proteins found exclusively in muscle nuclei from sedentary mice exhibit nuclear processes tied to transcription and chromatin remodeling, the nuclear proteome unique to exercised mice seems to indicate an emphasis on catabolic processes linked to the proteasome. It is important to note that these data cannot exclude the likelihood that contaminant cytoplasmic proteins were also included with isolated nuclei.

To gain deeper insights into the relative differences in the respective proteomes, we performed pathway analysis on the 1,672 proteins that were shared between both sedentary and exercise mice. We further categorized these proteins based on their fold change to highlight potential distinctions between sedentary and exercise nuclei. Using a fold change (FC) greater than 1 based on abundances detected by LC-MS, 32% (542 proteins) out of the 1,672 were more abundant in exercised mice, whereas 68% (1,130 proteins) were more abundant in sedentary mice. Nevertheless, the pathway analysis (FDR < 0.05) of these proteins indicated that sedentary mice (*orange*) demonstrated an abundance of post-transcriptional processing pathways, while exercised mice appeared to enrich pathways associated with translation and ribosomal machinery (*blue*) (**Figure 4F**).

In our pursuit of a more comprehensive understanding of how exercise impacts the nuclear proteome in skeletal muscle, we focused on the 1,672 proteins common between sedentary and exercised mice and specifically focused on proteins associated with nuclear transport and chromatin remodeling. Our pathway enrichment analysis (FDR < 0.05), unveiled a significant number of proteins linked to nucleocytoplasmic transport, with the largest number of proteins included in mRNA transport (**Figure 4G**). To delve deeper into the distribution of these proteins, protein abundances detected by LC-MS were used to calculate fold changes. Using a fold change greater than 1 (FC > 1) revealed that nearly 85% of the 67 proteins connected with mRNA transport were more abundant in sedentary mice compared to their exercise counterparts. Within this group, there was a notable change in various subunits of the nucleoporin complex (NUPS), which constitute the primary components of the nuclear pore complex – a key hub of nucleocytoplasmic transport. Additionally, proteins associated with the TREX multiprotein complex, specifically THO complex proteins (THOC2, THOC5, THOC7), known to interact with the nuclear pore complex coupled during mRNA export(31, 32), were also more prevalent in sedentary mice (**Supplemental Figure S2A**). Although a few proteins exhibited higher levels in exercise mice, the overall trend suggest a decrease in proteins involved in nucleocytoplasmic transport in response to acute exercise at 1-hour post-exercise.

Shifting our focus, we investigated proteins linked to histone and chromatin modifications shared between sedentary and exercise mice (**Figure 4H**). Over 200 proteins were enriched in pathways associated with chromatin organization and chromatin remodeling (FDR < 0.05). To disentangle the specific effects of exercise, we highlighted common histone markers indicative of chromatin remodeling with a FC > 1. This includes histone deacetylation (n = 19), acetylation (n = 19), ubiquitination (n = 6), and methylation (n = 6) (as shown in **Supplemental Figure S2B**). Notably, a substantial portion of proteins (50-60%) associated with histone deacetylation or histone acetylation as well as proteins connected with histone ubiquitination and methylation were higher in sedentary mice. We also identified proteins belonging to the SWI/SNF superfamily (n = 15), well-known for their role in influencing chromatin remodeling and accessibility(33, 34) (**Supplemental Figure S2B**). These included AT-rich interaction domain proteins (ARID1A, ARID1B, ARID2) and various SWI/SNF-related matrix-associated actin-dependent regulators of chromatin proteins (e.g., SMARCA2, SMARCA4, SMARCC2, SMARCE4). Much like our findings regarding histone-modifying proteins, constituents of the SWI/SNF superfamily were more abundant in sedentary mice than in exercise mice, hinting at their potentially reduced influence on nucleosome-DNA interactions post-exercise.

Collectively, these findings offer a novel and expanded perspective, shedding light on the influence of exercise on a broad array of proteins associated with transcriptional and post-transcriptional regulation, catabolic processes, translational control, nucleocytoplasmic transport, and chromatin remodeling, providing insights that were previously unexplored.

### Exercise modulates post-transcriptional processing in skeletal muscle nuclei

The statistical comparison between the nuclear proteomes of sedentary and exercised mice revealed notable differences. Specifically, we identified 224 proteins with significant changes (FDR < 0.1), with 135 proteins showing higher levels in the exercised group and 89 proteins displaying higher levels in the sedentary group (**Figure 5A**). As seen in supplemental Figures S3A and S3B there were sarcomeric and mitochondrial proteins in the 224 proteins with a significant change. There is a high likelihood that these proteins reflect cytoplasmic remnants that remained with the nuclear enrichment. Thus, we filtered all statistically significant proteins based on annotated nuclear subcellular localization. This refinement resulted in a total of 72 nuclear-enriched proteins (accounting for 32.1% of the 224 proteins) that displayed significant differences after exercise. These proteins were associated with biological processes like mRNA processing, RNA splicing, messenger ribonucleoprotein complex assembly, and mRNA transport (**Figure 5B**). Furthermore, KEGG pathway analysis highlighted enrichments in the spliceosome, ribosome, HIF-1 signaling, and ubiquitin-mediated proteolysis (**Figure 5C**). Notably, the proteins enriched in each of these processes and pathways generally exhibited higher abundance in sedentary mice compared to exercise mice, indicating a downregulation of these proteins in the nucleus either through degradation or transport following exercise. To support this, gene set enrichment analysis (exercise vs. sedentary) showed processes like RNA processing, RNA splicing, RNA 3’-end processing, and RNA transport were more enriched in sedentary mice. Nuclear-resident proteins of various compartments (e.g., Envelope, Nucleolus, Paraspeckles, Spliceosome) were also more enriched in sedentary mice, thus supporting the downregulation of nuclear proteins at 1-hour post-exercise (**Supplemental Figure S3C and S3D**).

**Figure 5.**
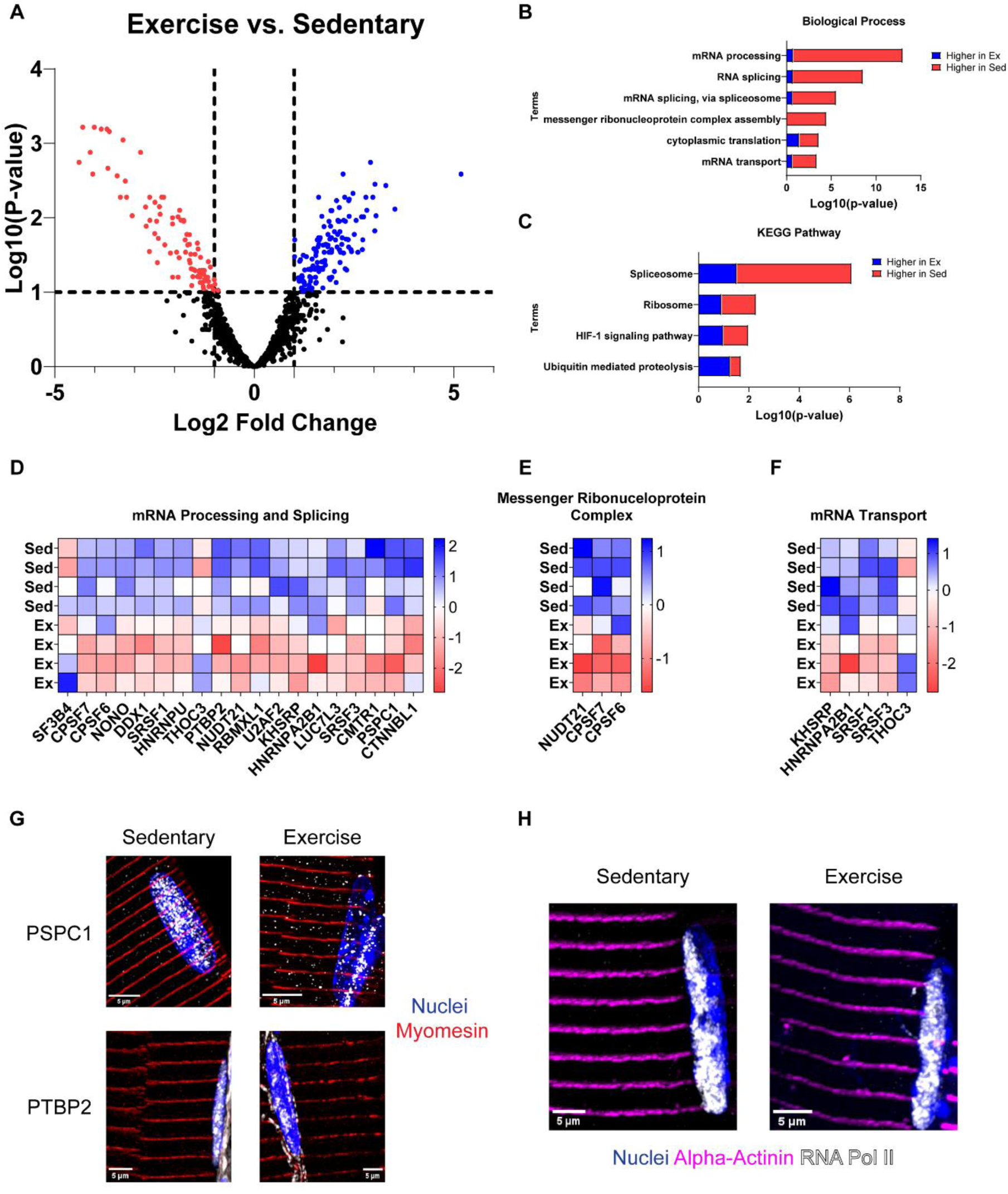
Statistical comparison between sedentary and exercise nuclear proteomes. **A)** Volcano plot of statistically significant (FDR < 0.1) proteins found in sedentary and exercise muscle nuclei. (*blue* is significantly higher after exercise; *red* is significantly lower after exercise). **B and C)** Significantly enriched (FDR < 0.05) biological processes and KEGG pathways of significant proteins. Proportional equivalents of the number of proteins within biological processes and KEGG pathways are shown that were either higher in exercise (*blue*) or higher in sedentary (*red*). **D-F)** Heatmaps of significant proteins of the enriched biological processes in panel B, including mRNA processing & splicing (**D**), messenger ribonucleoprotein complex (**E**), and mRNA transport (**F**). **G)** Images of skeletal muscle nuclei (DAPI; *blue*), z-line protein myomesin (*red*), and nuclear proteins paraspeckle protein component 1 (PSPC1) or polypyrimidine tract binding protein 2 (PTBP2) (*white*) in longitudinal sections of tibialis anterior muscle from sedentary (*left*) and exercise (*right*) mice. **H)** Images of skeletal muscle nuclei (DAPI; *blue*), z-line protein alpha-actinin (*magenta*), and nuclear-residing RNA polymerase II (RNA Pol II) (*white*) in longitudinal tibialis anterior muscle from sedentary (*left*) and exercise (*right*) mice. Scale Bar = 5 µm.

Specific biological processes, such as mRNA processing and splicing, are illustrated in heatmaps (**Figure 5D-F**), demonstrating an overall higher abundance of constituent proteins in sedentary mice. This includes proteins like U2 small nuclear ribonucleoprotein auxiliary factor 2 (U2AF2), serine/arginine-rich splicing factors 1 and 3 (SRSF1, SRSF3), beta-catenin like-1 (CTNNBL1), and Luc7-like-3 pre-mRNA splicing factor (LUC7L3). Many of these proteins serve multifaceted functions that overlap with other significantly enriched processes, such as cleavage and polyadenylation-specific factors 6 and 7 (CPSF6 and CPSF7, respectively), which are involved in 3’-end polyA signaling, essential for pre-mRNA maturation and mRNA export(35–38).

Heatmaps representing enriched KEGG pathways exhibited a similar trend, with the majority of proteins in each pathway being more abundant in sedentary mice, except for ubiquitin-mediated proteolysis (**Supplemental Figure S3E-H**). Proteins associated with ubiquitin-mediated proteolysis, including Ribosomal Protein S27a (RPS27A; Ubiquitin), Elongin B (ELOB), and Elongin C (ELOC), were found in higher quantities after exercise (**Supplemental Figure S3F**). While elongins are known to contribute to the transcription elongation process of pre-mRNA(39), they are also involved in the ubiquitination of target proteins as core components of cullin-RING-based ECS E3 ubiquitin-ligase complexes(40–42). The increased presence of elongins coupled with elevated ubiquitin levels after exercise lends support to the possibility of nuclear proteolysis post-exercise, in line with the indications of catabolic processes and proteasome involvement in our enrichment analysis (**Figure 4F**).

To validate some of these findings, we employed super-resolution microscopy on longitudinal sections of the tibialis anterior muscle (15) to visualize changes in nuclear proteins following exercise (**Figure 5G**). In our investigation, we focused on two proteins: paraspeckle component protein 1 (PSPC1) and polypyrimidine tract binding protein 2 (PTBP2) (Figure 5G *white*), both of which play crucial roles in nuclear-localized RNA processing. PSPC1 was readily observed in DAPI-stained nuclei within skeletal muscle of sedentary mice, appearing as concentrated clusters dispersed throughout the nucleoplasm. PTBP2 exhibited a more diffuse distribution throughout the nucleoplasm of sedentary mice, reflecting its extensive involvement in RNA binding. However, both PSPC1 and PTBP2 exhibited visually reduced content in skeletal muscle nuclei in response to exercise, aligning with the findings from our proteomic analysis. As seen in figure 5G, PSPC1 display greater speckling and dispersion within the cytoplasm of muscle fibers in exercise mice, whereas PTBP2 showed reductions in nuclear localization but remained on the outer perinuclear boundary. In contrast, the transcriptional multiprotein complex, RNA polymerase II, showed distinct localization in muscle nuclei with no changes after acute exercise (**Figure 5H**). Taken together, our data suggests that proteins associated with nuclear post-transcriptional processes appear to be reduced, we suggest through transport processes, within the nucleus one hour after acute exercise.

## Discussion

In the present study, our goals were to validate an approach using an enriched nuclear fraction from skeletal muscle for proteomics and to use this method to help define the skeletal muscle nuclear proteome in response to acute treadmill exercise. We found that our method resulted in significant enrichment of nuclear proteins including many not previously detected from whole muscle proteomics. Applying this approach to exercise, we successfully isolated skeletal muscle nuclei enhancing the identification of nuclear proteins including transcription factors. We determined that at 1-hour post-exercise, the predominant changes in nuclear proteins were linked to mRNA processing and splicing. These findings demonstrate an enhanced approach for probing nuclear protein changes in skeletal muscle in unbiased large-scale proteomic studies as well as highlighting changes to proteins linked to post-transcriptional processes after acute exercise.

The skeletal muscle nuclear proteome has been a major challenge in proteomic studies as the large and overabundant contractile proteins often obscure the detection of smaller, low abundant nuclear proteins. For example, the largest contractile protein titin comprises up to 16% of the total protein mass in whole skeletal muscle. With the addition of myosin, actin, myomesin, and others, 9 total proteins encompass over 50% of the total protein mass in whole skeletal muscle, thus limiting nuclear protein identification(8). Prior studies often utilize subcellular fractionation in combination with mass spectrometry(11, 12, 43–45) to improve the resolution of compartmental proteins(46–48). However, isolation of skeletal muscle nuclei has been historically challenging. Here, we successfully isolated an enriched sample of skeletal muscle nuclei using a modified fractionation approach to achieve deeper coverage of nuclear proteins compared to whole skeletal muscle.

Biochemical fractionation has been a long-standing experimental approach for the isolation of cellular compartments and their constituents(9, 49). In general, homogenized tissue is centrifuged over a density gradient for the physical separation of its compartments. We adapted such an approach(22) allowing for the separation of intact muscle nuclei while simultaneously depleting larger contractile proteins, enhancing our identification of nuclear classified proteins. Indeed, we show a greater proportion of nuclear proteins in isolated muscle nuclei compared to whole muscle that largely reflected transcription, rRNA processing, splicing and mRNA transport pathways. Additionally, isolated nuclei also enhanced identification of DNA binding proteins and transcription factors, many of which are well known in the muscle field. While alternative strategies involve isolating fluorescently tagged nuclei from fresh tissue, our data highlight the benefits of this approach including the use of frozen skeletal muscle, removing the need of transgenic models, as well as its potential application across species(50–52). Thus, these results define the successful isolation of skeletal muscle nuclei and its utility for the study of the nuclear proteome.

We then applied this approach to study alterations in the skeletal muscle nuclear proteome 1-hour after acute exercise. We found 224 proteins that were changed at 1-hour post-exercise with the most prominent effect on proteins linked to mRNA processing and splicing. Surprisingly, we found that mRNA processing and splicing factor proteins were decreased 1-hour post-exercise which are in contrast to studies showing their upregulation in the context of aging(13, 53). Ubaida-Mohien et al. found 57 proteins of the spliceosome complex were higher during aging in human skeletal muscle coupled with higher rates of alternatively spliced transcripts(53). However, data from the same group highlighted that spliceosome machinery and processing proteins were negatively affected with physical activity(54). Thus, in conjunction with our data 1-hour post-exercise, supports a paradigm in which exercise results in decreased nuclear localization of RNA splicing-associated proteins.

Moreover, some of the proteins we found linked to RNA processing and splicing like PSPC1, PTBP2, and NUDT21 highlight a potential impact of exercise on the RNA processing network. PSPC1 together with the protein NONO form RNA-protein structures known as paraspeckles(55–57). NONO also decreased 1-hour post-exercise, suggesting reduced RNA processing sites in the nucleus. Moreover, decreases in RNA processing proteins like PTBP2 and NUDT21 implicates the effects of exercise on splicing, as these proteins are often a part of multi-protein complexes recruiting other RNA processing factors(35, 37, 38, 58–60), several of which are also reduced 1-hour post-exercise (i.e., CPSF6 and 7). Thus, acute exercise results in decreased nuclear localization of proteins linked to the network of RNA processing and splicing proteins in skeletal muscle nuclei 1-hour after exercise. At this time point, we suggest that these changes in nuclear localization are downstream of nuclear transport, but more work is needed to define the mechanism. Importantly, these data provide insight into an unexplored area of exercise that contributes to our understanding in post-exercise responses and adaptations.

Immunohistochemistry in muscle sections to visualize localization changes after exercise showed greater dispersion of PSPC1 in the cytoplasm whereas PTBP2 was restricted to the perinuclear boundary. To our knowledge, these are the first visual evidence of these nuclear proteins in skeletal muscle and future studies might explore nuclear transport mechanisms that might be responsible for decreases in nuclear proteins post-exercise. Alternatively, we cannot dismiss the possibility of nuclear protein turnover as subunits of the proteasome were exclusively found in nuclei after exercise. Nuclear localization of the proteasome has been shown in other cell types(61–64) however its involvement in skeletal muscle nuclei has not been explored. Additionally, nuclear proteins were only measured at 1-hour post-exercise and in sedentary controls, thus warranting analysis of additional post-exercise timepoints to uncover the dynamic patterns of nuclear protein changes after exercise. Nonetheless, our data show that acute exercise leads to a prominent decrease in proteins linked to mRNA processing and splicing in skeletal muscle nuclei 1-hour after exercise.

In summary, our modified approach for the isolation of skeletal muscle nuclei enhanced the identification of nuclear proteins compared to whole muscle and provides a methodology to more deeply probe the skeletal muscle nuclear proteome with exercise. While we identified proteins in isolated nuclei related to RNA processing, RNA transport, and chromatin maintenance, the most prominent exercise effect of was on mRNA processing and splicing proteins 1-hour after exercise. We suggest that these methods can be used to more deeply asses nuclear protein changes in skeletal muscle and we highlight mRNA processing and nuclear transport as new target areas for research in exercise science.

## Data Availability

The data that support the findings of this study are available on request from the corresponding author.

## Supplemental Material

Supplemental Figures S1 – S3:

## Acknowledgements

None.

## MoTrPAC Study Group

**Primary Authors**

Ryan A. Martin, Ph.D.*^1,2^*, Karyn A. Esser, Ph.D.*^1,2^*

**Preclinical Animal Study Sites**

Ryan A. Martin, Ph.D.*^1,2^*, Mark R. Viggars, Ph.D.*^1,2^*, Collin M. Douglas, Ph.D.*^1,2^*, Karyn A. Esser, Ph.D.*^1,2^*

**Chemical Analysis Sites**

James A. Sanford, Ph.D.*^3^*, Zane W. Taylor, Ph.D.^3^, Joshua R. Hansen, M.S.*^3^*, Geremy C. Clair, Ph.D.^3^, Joshua N. Adkins, Ph.D.*^3,4^*

^1^*Department of Physiology and Aging, University of Florida, Gainesville, FL, USA, ^2^Myology Institute, University of Florida, Gainesville, FL, USA, ^3^Pacific Northwest National Laboratory, Richland, WA, USA, ^4^Department of Biomedical Engineering, Oregon Health and Science University, Portland, OR, USA*

The authors would also like to acknowledge Jonathan Bird for accessibility and contribution to the images from super resolution microscopy.

## Author Contributions

R.A.M., M.R.V., G.C.C., and K.A.E. conceived and designed research, R.A.M., M.R.V., C.M.D., J.A.S., Z.W.T., J.R.H., G.C.C., and J.N.A. performed experiments, R.A.M., M.R.V., J.A.S., J.R.H., G.C.C., and J.N.A. analyzed data, R.A.M., M.R.V., J.A.S., G.C.C., C.M.D., and K.A.E. interpreted results of experiments, R.A.M., M.R.V., and C.M.D. prepared figures, R.A.M., M.R.V., C.M.D., and K.A.E. drafted manuscript, R.A.M., M.R.V., C.M.D., J.A.S., G.C.C., Z.W.T., J.R.H., J.N.A. and K.A.E. edited and revised manuscript, R.A.M., M.R.V., J.A.S., J.N.A., G.C.C., Z.W.T., J.R.H., C.M.D., and K.A.E. approved final version of manuscript.

---

**Supplemental Figure S1.**
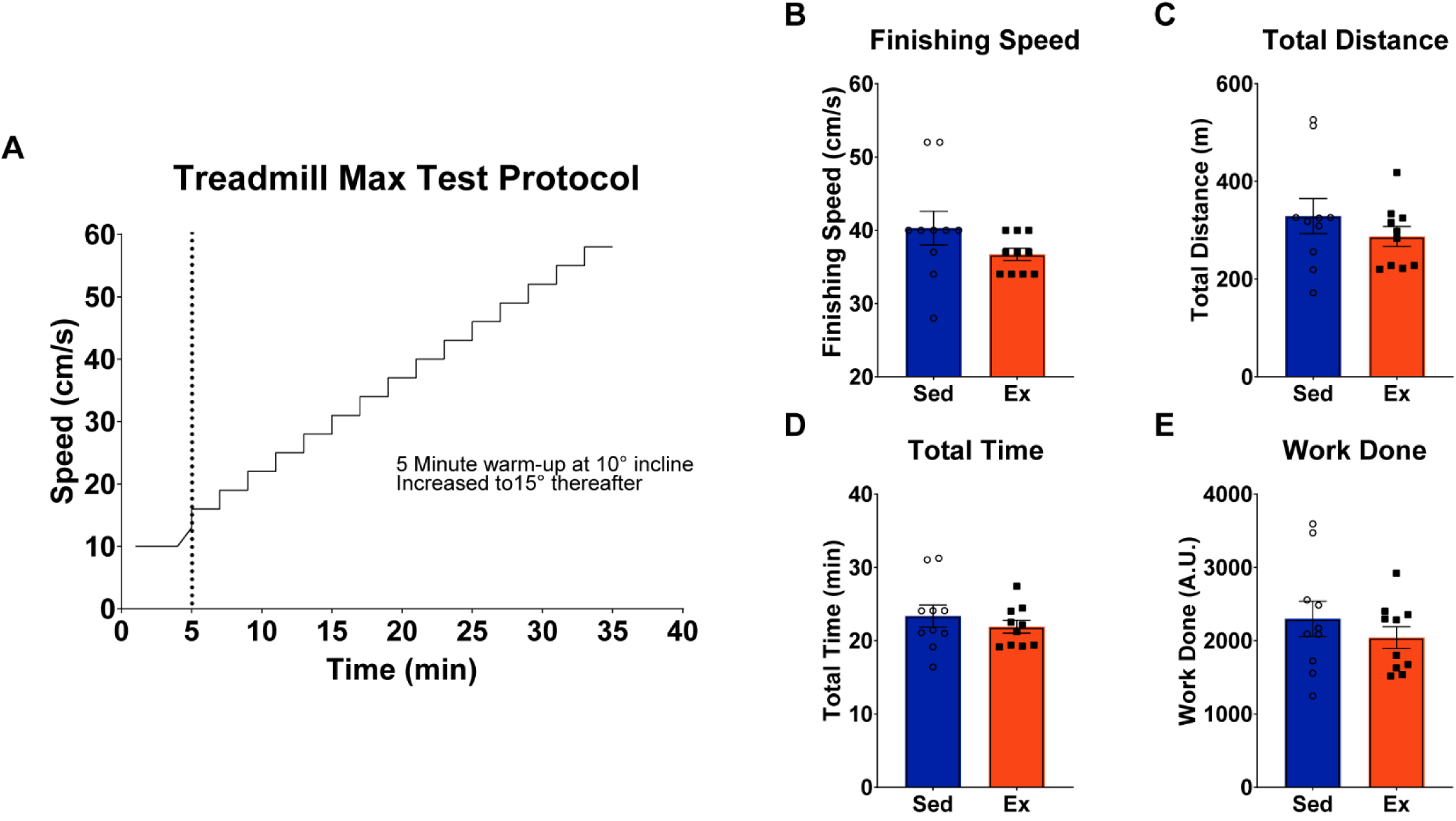
Parameters and performance of maximal treadmill exercise test. **A)** Protocol for the maximal treadmill test with a 5 minutes warm-up at 10° incline, increased to 15° incline with a progressive increase of 3 cm/s every 2 minutes until exhaustion. **B-E)** Maximal treadmill test performance measures of sedentary and exercise group mice including finishing speed (B), total distance (C), total time (D), and work done (E). Independent t-tests between Sed and Ex groups; all measures were not significant (NS; P > 0.05).

**Supplemental Figure S2.**
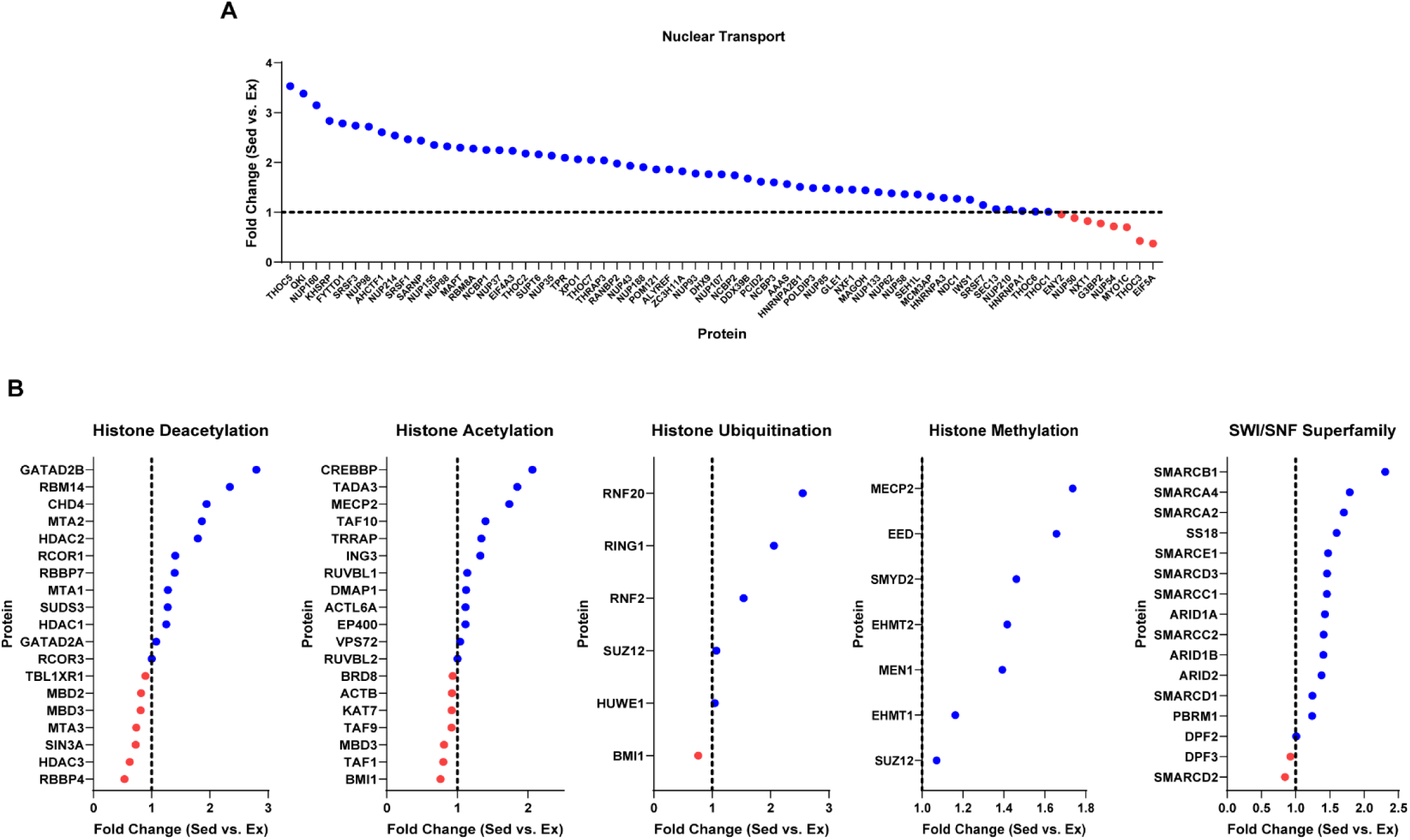
Nuclear transport and chromatin modifying proteins greater than 1-fold change in sedentary mice compared to exercise mice. **A)** Fold changes (Sed vs. Ex) of proteins enriched in mRNA transport pathway (n = 67). **B)** Fold changes (Sed vs. Ex) of enriched proteins related to post-translational modifications of histone methylation, histone acetylation, histone deacetylation, and members of the SWI/SNF superfamily.

**Supplemental Figure S3.**
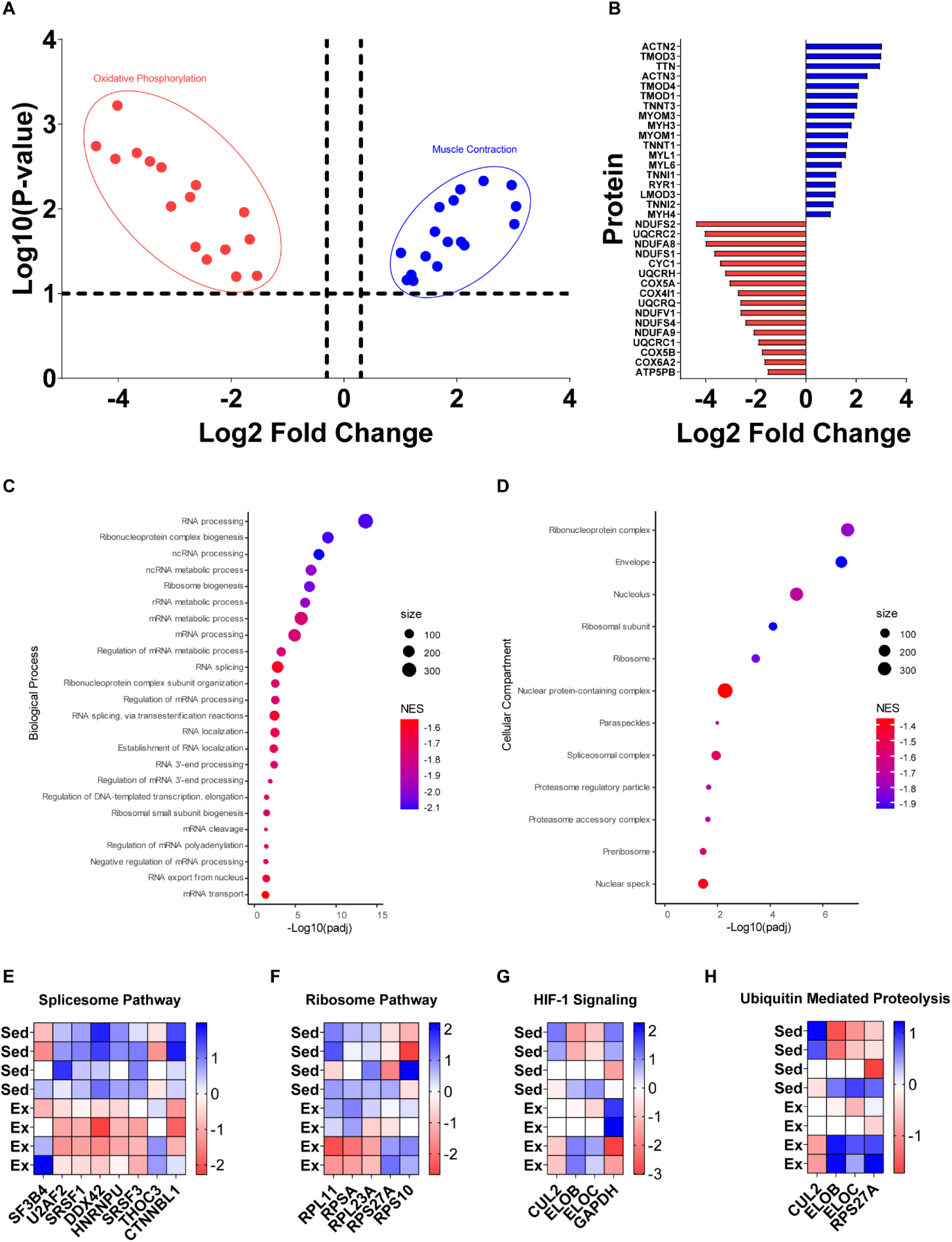
Gene Set Enrichment Analysis and KEGG pathways of significant proteins between sedentary and exercise mice. **A)** Volcano plot of proteins associated with either mitochondrial-associated (*red*) or muscle contraction (*blue*). **B)** Bar plot of proteins in (A) with respective log2 fold change related to mitochondria (*red*) or muscle contraction (*blue*). **C and D)** GSEA dot plots of Biological Process (C) and Cellular Compartment (D) of proteins (exercise vs. sedentary) nuclei. FDR < 0.05; NES = Normalized Enrichment Score; Size = number of proteins identified in pathway. **E-H)** Heatmaps of significant proteins of the enriched KEGG pathways in Figure 4 panel C, including spliceosome (E), ribosome (F), HIF-1 signaling (G), and ubiquitin mediated proteolysis (H).

